## Supplementary figures and images for "Cell type-specific network analysis in Diversity Outbred mice identifies genes potentially responsible for human bone mineral density GWAS associations"

### Figure 1-figure supplement 1

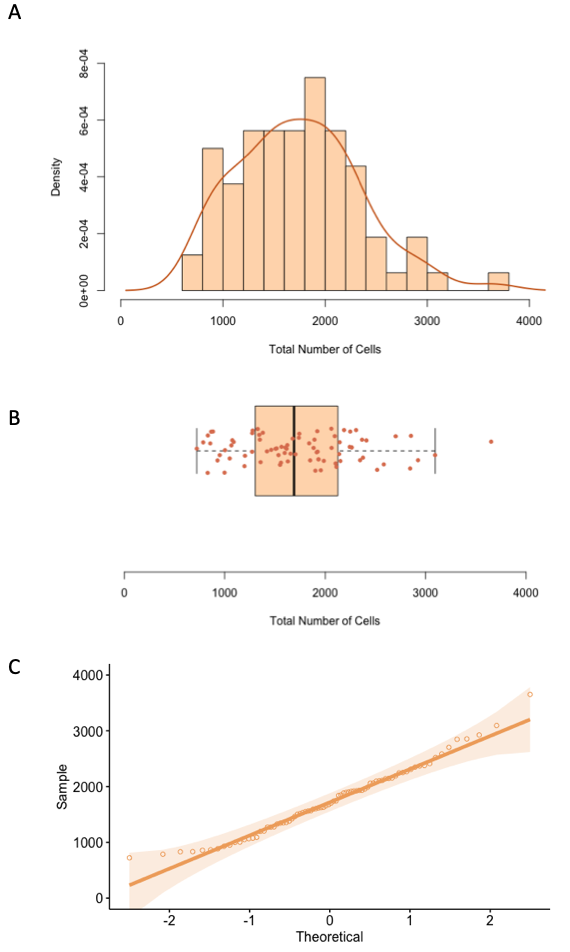

### Figure 2-figure supplement 1

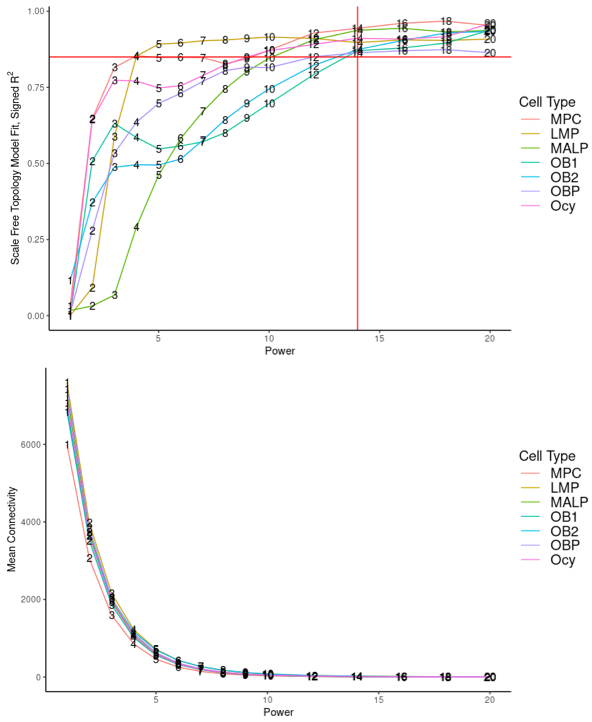

### Figure 4-figure supplement 1

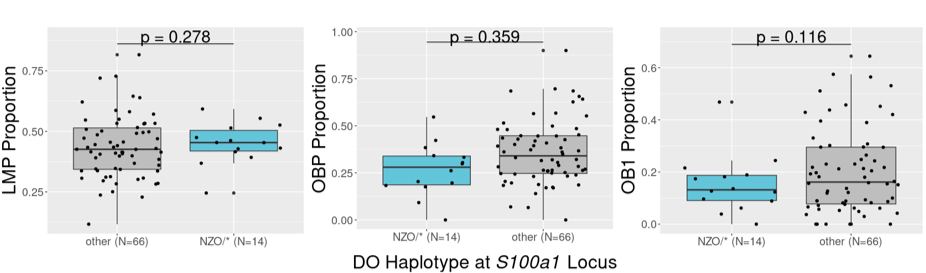
